# MEA Viewer: a High-performance Interactive Application for Visualizing Electrophysiological Data

**DOI:** 10.1101/146381

**Authors:** Daniel C. Bridges, Kenneth R. Tovar, Bian Wu, Paul K. Hansma, Kenneth S. Kosik

## Abstract

Multi-electrode arrays (MEAs) have been used for many years to measure electrical activity in ensembles of many hundreds of neurons, and are used in research areas as diverse as neuronal connectivity and drug discovery. A high sampling frequency is required to adequately capture action potentials, also known as spikes, the primary electrical event associated with neuronal activity, and the resulting raw data files are large and difficult to visualize with traditional plotting tools. Many common approaches to deal with this issue, such as extracting spikes times and solely performing spike train analysis, significantly reduce data dimensionality. Unbiased data exploration benefits from the use of tools that minimize data transforms and such tools enable the development of heuristic perspective from data prior to any subsequent processing. Here we introduce MEA Viewer, a high-performance interactive application for the direct visualization of multi-channel electrophysiological data. MEA Viewer provides many high-performance visualizations of electrophysiological data, including an easily navigable overview of all recorded extracellular signals overlaid with spike timestamp data and an interactive raster plot. Beyond the fundamental data displays, MEA Viewer can signal average and spatially overlay the extent of action potential propagation within single neurons. This view extracts information below the spike detection threshold to directly visualize the propagation of action potentials across the plane of the MEA. This entirely new method of using MEAs opens up new and novel research applications for medium density arrays. MEA Viewer is licensed under the General Public License version 3, GPLv3, and is available at http://github.com/dbridges/mea-tools.

## Introduction

Multi-electrode arrays (MEAs) are commonly used to record extracellular action potentials (eAPs) from in vitro and in vivo neural networks (Obien et al. 2014). Because action potential widths can range from about 0.2 to 4 milliseconds, depending on the neuron type (Bean et al., 2007) the sampling frequency required to reliably capture eAPs is commonly in the range of 10 to 25 kHz. Thus when using arrays with 120 electrodes, experiments can generate data at a rate of over 300 MB/min, resulting in large unwieldy files. It is common at this point to extract eAP information by spike detection and subsequent sorting, steps that simplify the task of data handling but also dramatically reduce data dimensionality. However, spike detection and sorting steps themselves are notoriously error prone (Rey et al., 2015) and information present in the lower frequency domains of analog recordings is not captured by spike detection. Methods and tools that routinely evaluate the performance of spike detection and spike sorting performance in an unbiased way can help avoid systematic errors by quickly identifying detection and sorting failures and reveal features in the raw data that may be difficult to access in sorted data.

Here we introduce MEA Viewer, a software package for high-performance visualization of multi-channel electrophysiological data recorded by a multi-electrode array system. MEA Viewer fills the void for tools to examine unprocessed data and allows direct inspection of the recorded extracellular signals superimposed with the results of spike detection and spike sorting and thus is ideal for examining the results of spike sorting routines. The primary audience for MEA Viewer is for end users of MEA systems who want to explore their data prior to doing heavy statistical analysis. MEA Viewer provides five main visualization interfaces: (i) the grid view displays the complete set of recorded extracellular signals, (ii) the signal comparison view allows users to select and display multiple recorded channels with superimposed spike data, (iii) an interactive raster view displays spike timestamp data for all recorded channels, (iv) the flashing spike view presents a spatial-temporal representation of the spiking behavior of the recorded channels and (v) the propagation signal view overlays aggregated spiking events, revealing an individual neuron’s multi-electrode propagation signal, as well as electrophysiological features typically masked by noise. We recently showed that MEAs can be used to monitor action potential propagation in single neurons (Tovar et al., 2017). Assigning spikes to a particular neuron is not typically possible with extracellular recording. However, by making these propagation signals easy to find, MEA Viewer makes it possible for the first time to unambiguously identify and explore the spiking behavior from single identified neurons. With MEA Viewer these spikes can be signal averaged to reveal electrodes with sub-threshold events among all MEA channels. Retaining these previously discarded events enables commonly used medium-density arrays (100-200 um electrode pitch) to reveal intracellular signal propagation in a high-throughput way, over multiple days. This may be especially useful in monitoring changes due to degeneration and in disease models.

## Design and Implementation

MEA Viewer was written in Python 3 and OpenGL Shading Language, making extensive use of the Python scientific stack (numpy, scipy, pandas, and h5py). The high performance of MEA Viewer is achieved by transferring a majority of the data processing to the graphics processing unit (GPU). Most of the interactions with the GPU are done with vispy (Campagnola et al. 2015), a relatively high-level Python interface to modern graphics hardware through the use of OpenGL. The rest of the user interface was created using PyQt and the Qt GUI library, enabling the application to operate across platforms (currently tested on Windows 7 and Mac OS X 10.10+).

MEA Viewer was written to be a general-purpose electrophysiology display application for large (0.5-1.5 GB) multi-stream data files. It accepts input data given in the form of Hierarchical Data Format version 5 (HDF5) files for extracellular signal recordings and comma-separated value (CSV) files for spike timestamps. HDF5 is an open file format stewarded by the HDF Group, and is the file format adopted by the Neuroscience without Borders initiative (Teeters et al. 2015). Other projects such as NEO (Garcia 2014), G-Node (Sobolev 2014), or NSDF (Ray 2016) seek to provide libraries to read electrophysiological data from a variety of formats, but without there being a clear winner among the three we chose to focus our efforts on supporting only HDF5 files. Spike timestamp data can be generated using tools available with MEA Viewer, or from any other spike detection and sorting program, then converted to a CSV file compatible with MEA Viewer (see supplementary materials). MEA Viewer seeks to remain agnostic to a chosen spike sorting routine, thus due to the vast number of options in spike sorting, and with no clear sorting method or implementation widely adopted, a common file format like CSV was needed to provide for maximum interoperability. MEA Viewer is currently designed to interoperate with Multi Channel Systems’ 120 electrode count MEA data, converted to HDF5 using Multi Channel Systems’ Data Manager conversion utility however other data formats are straightforward to add. A brief overview of the data-conditioning steps required is shown in Fig. 2A.

**Fig. 1.**
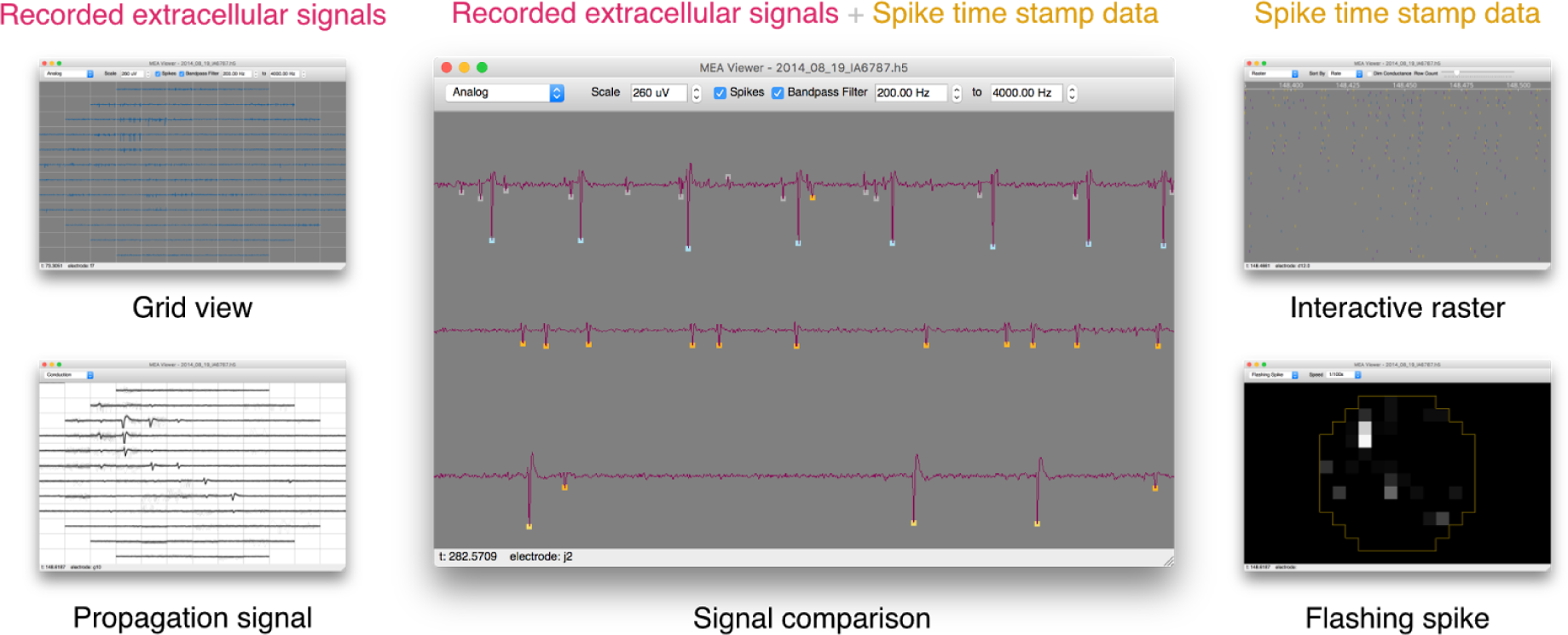
MEA Viewer provides 5 interactive visualizations of electrophysiological data. Recorded extracellular signals (top left) and spike timestamp data (top right) can be overlaid (middle) for unbiased investigation of MEA data. Multiple views are provided to easily navigate the entirety of the recorded signals, providing the ability to probe spike timing relations (bottom left), view and identify signals consistent with axonal propagation (bottom left), and verify spike detection and sorting performance (middle).

**Fig. 2.**
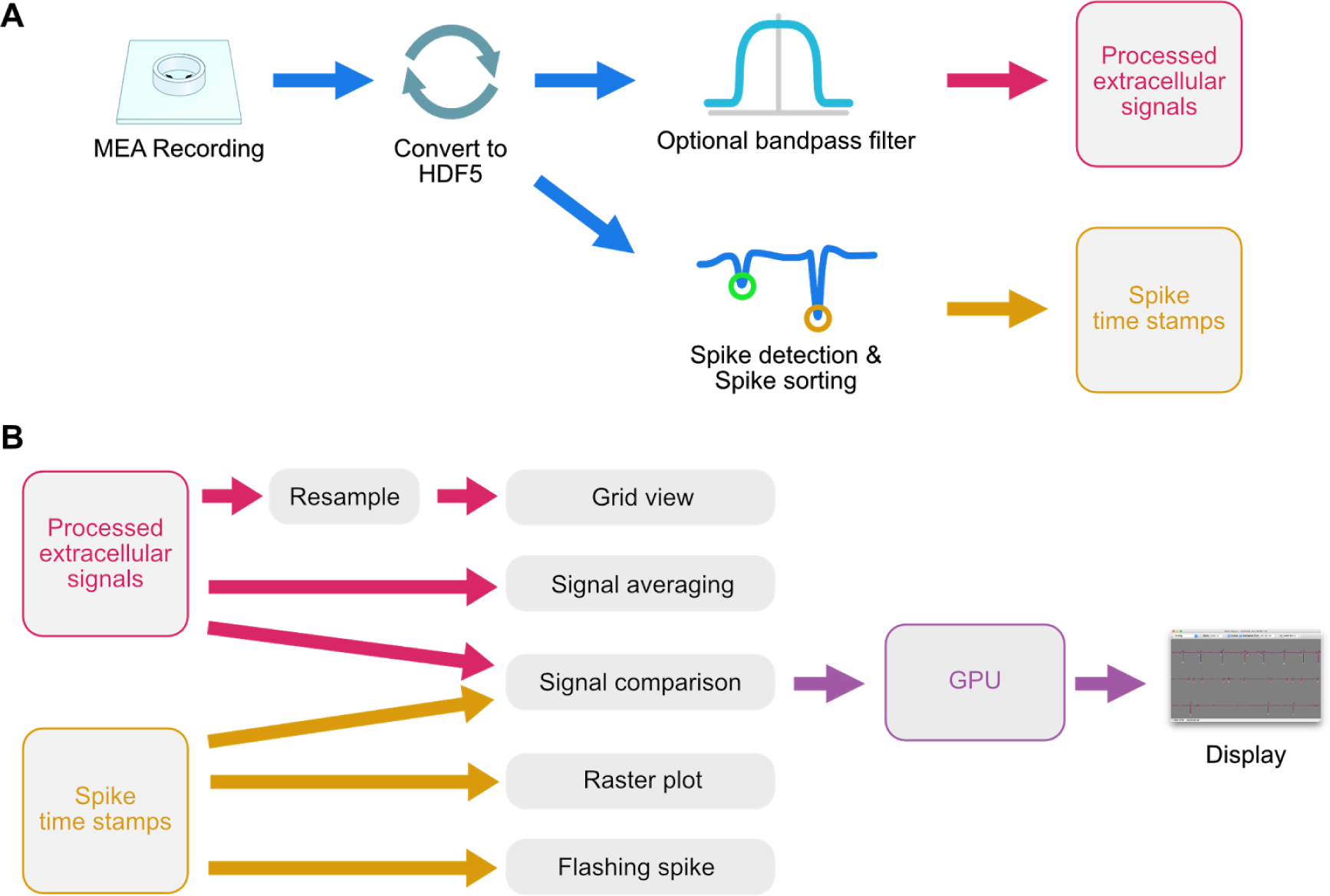
Pre-processing and display workflow used by MEA Viewer. (A) Extracellular signals are recorded with a commercial MEA system, then converted to a Hierarchical Data Format (HDF5). Data can then be optionally filtered and processed by a spike detection and spike sorting engine. These steps occur outside of MEA Viewer, but can be completed with tools in the overall mea-tools package. (B) The recorded extracellular signals and spike time stamp data are imported into MEA Viewer and loaded into memory. Due to the large number of data points, recorded signals are resampled prior to display in the grid view. Once a user selects their desired visualization, MEA Viewer generates vertex data that is sent to the GPU of the host computer, then displayed.

Once input data is properly formatted for reading by MEA Viewer, GPU vertex data is created by the currently active visualization and sent to the GPU for rendering (Fig. 2B). Most visualizations create vertex data for the entirety of the dataset displayed and transfer it once to a vertex buffer object on the GPU. To allow for panning and zooming a transformation matrix is then updated each animation frame, eliminating the massive transfer of vertex data on a frame-by-frame basis. One notable exception to this methodology is with the grid view which displays the entirety of the originally recorded extracellular signals. Because the grid view often displays overviews of a gigabyte or more of analog data, it is impractical to send all of this to the GPU at one time due to memory limits on consumer grade GPU cards. Instead the analog data is resampled and updated during mouse drag and scroll events to only transfer the selected data. Typically data is downsampled for display as the number of data points encompassing the record is much larger than the number of pixels used to display it. When dealing with long-duration recordings of spiking neurons it is important to downsample in a way that preserves an accurate view of these relatively sparse events. Transferring every n^th^ point makes it unlikely for these points to land on spikes—creating a visualization that underestimates the actual spiking behavior. We sidestep this by using a simple method to downsample by calculating the number of pixels used to display each waveform (n_p_), then binning the waveform data into n_p_ bins. The minimum and maximum value for each bin are calculated and those points are sent to the GPU as vertex data, which is then displayed as an OpenGL line strip. This technique preserves an accurate view of the spiking behavior and reduces the total amount of data being transferred to manageable levels, preserving a fluid interaction with the data.

Many of the visualizations display data in a layout that mimics the geometry of the MEA. Arbitrary layouts can be specified in the software by subclassing the abstract Layout class and implementing the methods `coordinates_for_electrode` and `electrode_for_coordinates`, as well as providing `rows` and `columns` attributes specifying the number of rows and columns in the layout. These functions are mostly self explanatory, with all coordinate values given in (column, row) tuples and electrode values given as strings.

The user interface of MEA Viewer makes it easy to switch between any of the visualizations while maintaining a consistent position in the data record. For instance, after panning to a specific part of the data in the interactive raster view, switching to the flashing spike view will show data from that same time point. This makes it easy to coherently navigate the data record and rapidly switch between views of the original extracellular recordings and views of the spike timestamp data.

## Results

The entry point for displaying extracellular signals with MEA Viewer is the grid view. From this window, users can view the recorded analog data from all electrodes, easily pan forward and reverse in the time record and zoom in and out at chosen locations in time and signal amplitudes. Specific channels can then be selected and a comparison view can be activated to display recorded signals from a subset of the channels overlaid with spike detection and sorting data (Fig. 3A). Detected spikes are color-coded by markers indicating sorted spike groups. Spikes detected on multiple electrodes but originating from single neurons are indicated by gray markers. Navigating the data in this way allows for unbiased assessment of spike detection and sorting performance of any data record, and enables the rapid visualization of redundant spike information, which can be underappreciated when analyzing MEA data (Tovar et al., 2017). The obvious utility of this visualization in MEA Viewer is that common spike-detection and -sorting routines can be easily assessed in this view (Fig. 3B). As detection and sorting errors can have large effects on measured spike rates, MEA Viewer can be used to spot cases like these early in the data analysis process.

**Fig. 3.**
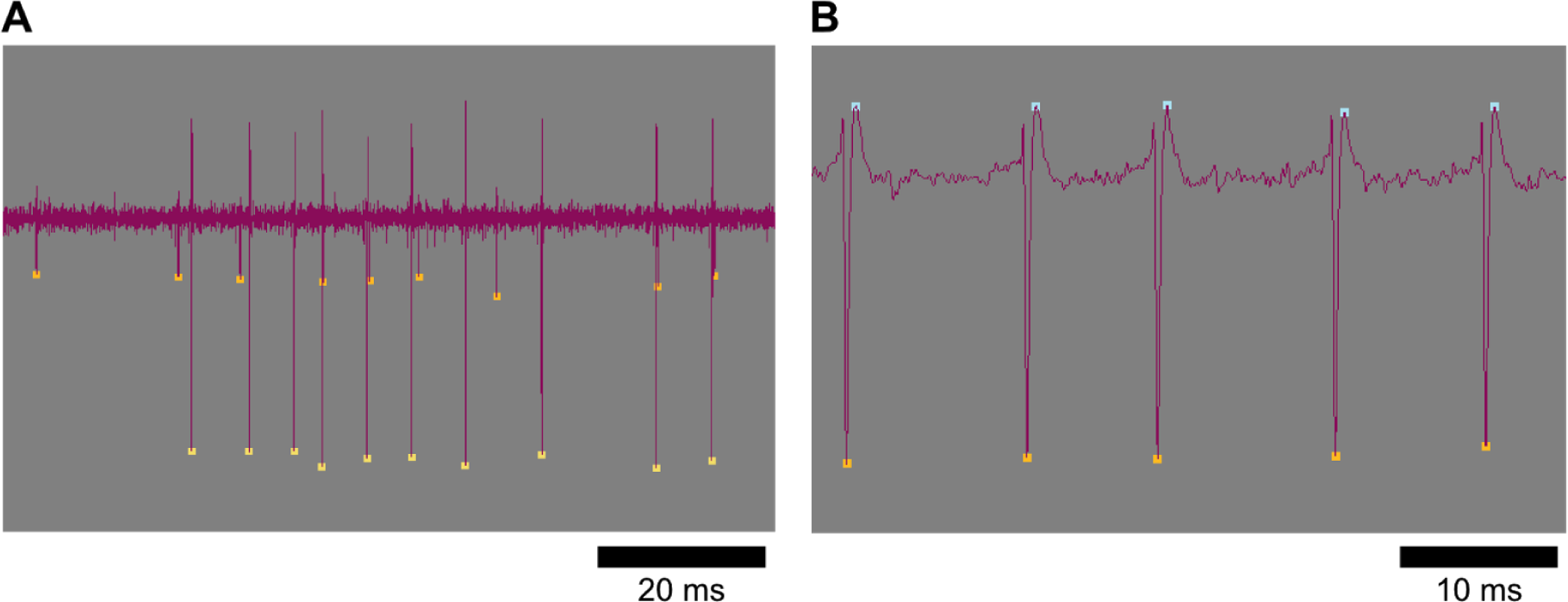
A signal comparison view displays recorded extracellular signals with overlaid spike timestamp data. (A) The signal comparison view allows for the inspection of spike detection and spike sorting accuracy. Here, two distinct waveforms are present and correctly sorted (see orange and yellow markers on small and large amplitude spikes, respectively). (B) An example showing a systematic error in spike detection on this large-amplitude signal. A false positive spike detection (blue) consistently occurs during the re-polarization phase of the action potential. Errors in spike processing like these are hard to detect without comparisons to the raw original recording and can have large effects on computed spike train statistics.

An additional unique feature of MEA Viewer is that it easily allows visualization of instances where the acquisition system captures action potential propagation in single neurons. Recent work with cultured neurons from mouse or human induced pluripotent stem cells (iPSCs) demonstrates that signal propagation within individual neurons can be monitored by detection of near-coincident spikes among MEA electrodes represent (Tovar et al. 2017). Because they arise from single neurons, these signals can be aligned and averaged to reveal the extent of signal propagation within one cell within a neural network, immediately giving a display of the spatial extent of signal propagation. The propagation signal view in MEA Viewer overlays these aggregated spike waveforms to visualize a single neuron’s propagation signal across an array, as well as other associated events occurring close in time to spikes in identified cells. We routinely record action potential propagation on multiple electrodes of arrays with inter-electrode distances of 100-200 um.

MEA Viewer signal averages spikes from single neurons by comparing event times of two channels when they show stereotyped coincident spiking (defined by many repeated events within 0.7 ms of each other). The first is the ‘reference’ channel and the second is the ‘test’ channel. The reference channel can alternatively be chosen by extracting spike times from single sorted channels. Once a list of event times is chosen using either method, a 20 ms window of data is extracted from every data channel centered on each spike time in the reference channel (Fig. 4A). The averaged waveform for each data channel are then superimposed with individual waveforms, and the waveforms from all data channels are displayed in a layout mirroring the MEA geometry. Signal averaging hundreds of spikes in this way often reveals electrodes throughout the array with sub-threshold events and gives an overview of behavior at all electrodes time-locked to the spike events occurring in the reference channel (Fig. 4C). Each channel showing highly correlated events likely reflects action potential propagation within a single neuron measured at different locations. In this example, the earliest action potential is in the top left and propagates to the bottom right (Fig. 4B).

**Fig. 4.**
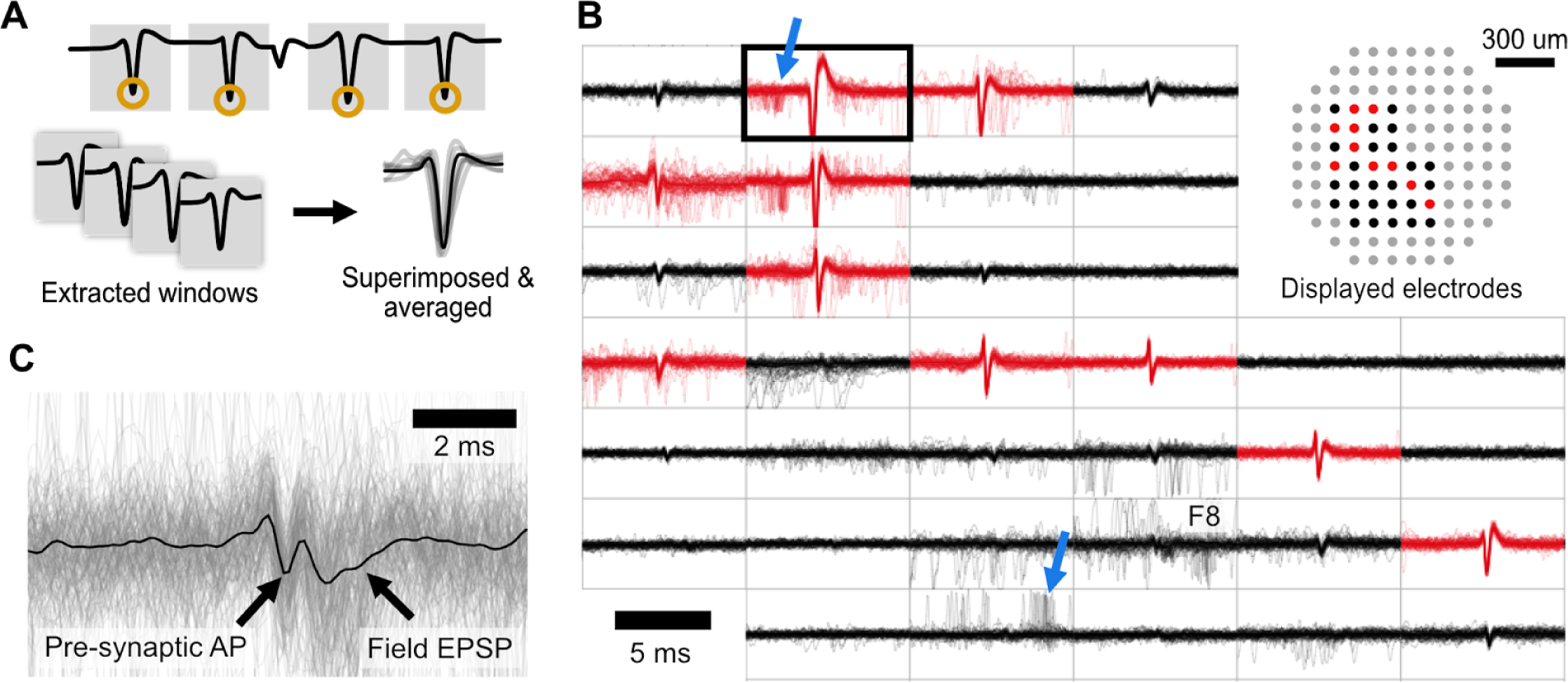
The propagation signal view overlays aggregated waveforms from all channels during repeated spike events from a reference channel, revealing axonal propagation and coupled pre- and post-synaptic spiking. (A) The reference channel (black outline) can be chosen as a single spike-sorted channel or two coincident channels. For each spike time in the reference channel, a 20 ms window is extracted from all other channels. The collected windowed waveforms for each channel are superimposed and averaged providing an overview of time-locked events across the array. (B) Example traces from cultured mouse hippocampal neurons grown on a 120 electrode count MEA (100 µm inter-electrode spacing), with the reference alignment channel outlined in bold. Channels shown in red are above spike detection threshold. In this example, there are numerous channels exhibiting time-locked spiking with the reference channel (well-defined central spikes, in red). These events are due to action potential propagation down an axon, originating from the top left corner and propagating to the bottom right. A cloud of spikes preceding (top blue arrow) and following (bottom blue arrow) the propagation signal are also visible, consistent with expectations for presynaptic and postsynaptic partners of the reference neuron, respectively. Relationships such as these revealed by MEA Viewer can be subsequently confirmed experimentally. (C) Enlarged and rescaled view of electrode labeled F8 in (B). Signal averaging reclaims features typically masked by noise, in this instance a waveform consistent with a excitatory postsynaptic potential (EPSP) and possibly the presynaptic AP are revealed.

The propagation signal view also reveals additional network features that are time-locked to spiking in the propagation signal. For example, in the example of the propagation signal shown (Figure 4B) there are spikes from other units that occur with timing expected from neurons with direct pre- or postsynaptic coupling to the neuron giving rise to the propagation signal. These events are distinguished from propagation signals by their longer time delay, typically between 1 and 10 ms, and their increased jitter with respect to the reference electrode, shown by the cloud of spikes seen in the overlaid waveforms (Figure 4B, blue arrows). In contrast, waveforms attributed to single neurons across multiple electrodes have very little jitter, and superimpose well over multiple instances (Tovar et al., 2017). Averaging spikes in the reference channel of a propagation signal also unmasks features typically obscured by noise, for instance a signal consistent with the properties expected from excitatory post-synaptic potentials (EPSP; Fig. 4C), something normally not detected in by extracellular recording of cultured neurons.

The interactive raster view in MEA Viewer displays spike time data from all unprocessed or spike-sorted data channels. Raster plots easily show array-wide activity patterns but are static displays. A static raster plot displays 3 minutes of data at a resolution of ~130 ms/px on a 1400 pixel monitor. This is in comparison to the high temporal resolution provided by MEAs (typically 50 µs or better) and spiking behavior that can exceed 20 Hz during high-frequency bursts. MEA Viewer can display spike dynamics at user-chosen time scales, sidestepping the limitations of raster plots. MEA Viewer also offers many channel-ordering schemes which define a channel’s row position to facilitate exploratory investigation. To provide a clearer view of array spiking dynamics, data in the interactive raster view can be ordered by activity level, with the most active channels on top, by their spike latency from an arbitrary time or by a fixed electrode order. Lastly, the interactive raster view has the option of dimming redundant spikes attributed to propagation signals. This feature clearly reveals the level of extraneous information present that should be accounted for prior to subsequent spike train analysis.

Finally, the flashing spike view displays the spatial-temporal patterns of spike activity. Array activity is animated by flashing a marker at a channel’s physical position when a spike arrives at that channel. Playback time is adjustable (from real time to 1/1600 of real time), to observe the dynamics among active channels. As subsequent spikes arrive, the spatial and temporal characteristics present are obvious. This view is meant to make obvious the spatial and temporal characteristics present in array-wide data as well as demonstrating the very short spike latency among electrode components of a propagation signal compared to array wide spiking (video available in Supplementary Materials). The flashing spike view is useful to assess data from experiments in which temporal spiking patterns are important to superimpose on spatial electrode profiles (Feinerman et al., 2008).

## Discussion

MEA Viewer was written to give users an interactive path towards exploratory data analysis. MEA Viewer supports rapid initial investigation of recorded electrophysiological data and gives users the ability to display the dynamics of the system under study prior to more sophisticated statistical analysis. Software packages such as the Matlab toolbox MEA-Tools (Egert et al. 2002) or the commercial NeuroExplorer focus heavily on statistical analysis of spike timestamp data without providing views of the analog data from all electrodes simultaneously. The open-source visualization tool, NeuroScope (Hazan et al. 2006), displays continuous time signals, but does not offer the performance of MEA Viewer to quickly navigate large data sets. Software provided by MultiChannel Systems only replays recorded data from all channels in a single playback direction and does not have the ability to easily compare user-chosen channels, or to zoom and pan through the dataset. No other applications support visualization of signal-averaged propagation signals. These unique capabilities of MEA Viewer facilitate novel investigations of MEA data, as well as improve the speed and understanding of current analysis techniques.

The multiple visualizations in MEA Viewer present data in an unprocessed and unbiased way, and permit other less spike timing-centric analyses. Exploratory data visualization often identifies errors early on in analysis workflow, and is therefore vital before subsequent processing. Having a variety of complementary data visualizations enhances the user’s intuition of the system under study and allows for rapid hypothesis generation and more robust testing. Furthermore, tools that allow end users to approach MEA data beyond traditional spike train analysis provide novel opportunities for investigating diseases or compounds that affect axonal propagation or ion channel distribution. In addition, inquiries into developmental progression of axon excitability, especially useful when studying neuronal iPSC lines, are possible. Thus, MEA Viewer provides an attractive first entry point to all types of MEA data for examining the entire topology of features present in extracellular recordings, from traditional spike train analysis to observation and identification of single neuron propagation signals.

Measurement of action potential propagation has been previously reported, but with high-density arrays (Bakkum et al. 2013) composed of thousands of electrodes, or by more technically challenging dual electrodes (Khaliq and Raman 2005). In contrast, the visualization algorithms in MEA Viewer allow for the routine identification and measurement of these propagation signals in a non-invasive, high-throughput fashion over many days. Once MEA Viewer identifies propagation signals, subsequent analysis (e.g., developmental changes of electrophysiological properties), can occur in any analysis software of choice.

Because of its general applicability MEA Viewer is an attractive platform on which to build additional features. In its current state, MEA Viewer is primarily intended for exploratory data visualization. This is useful because it easily allows the large data sets that are produced by MEA recordings to be examined prior to subsequent statistical approaches. Future visualizations in MEA Viewer will focus on displaying data interpretation alongside the currently implemented views. The application of network modeling techniques to spike train data is one area that could benefit substantially from a simplified user interface, enabling more widespread adoption of these sophisticated techniques. Dynamic modeling approaches which produce functional networks, like the kinetic Ising model and generalized linear model (Roudi et al. 2015; Stevenson et al. 2008; Latimer et al. 2014), are powerful techniques for spike train analysis, but are not easily accessible. MEA Viewer could be extended to provide an intuitive interface to refine input parameters of various network models, and display an attractive view of the resulting network. Users would then be able to perturb the generated functional model - for example, by removing individual neurons, groups of nodes, or specific connections within the network - and see the resulting output in the network display. Additional features such as these would only improve the usefulness of the already implemented core functionality of MEA Viewer.

## Availability

MEA Viewer is available on Github at https://github.com/dbridges/mea-tools. Links to precompiled Windows binaries are also available at the same address. MEA Viewer is licensed under the GPLv3. User suggestions and contributions are welcome.

## Methods

### Cell culture

All animal procedures were approved by the University of California’s institutional IACUC protocol, in accordance with NIH policy. Neurons were prepared from C57BL/6 mice. Briefly, hippocampi were dissected from postnatal day 1 mice, dissociated using papain and trituration, and plated at a density of 550 cells/mm^2^ on a confluent bed of glia. Cells were incubated at 37 °C with 5% CO_2_ and maintained in MEM supplemented with 5% heat-inactivated fetal calf serum and Mito+ serum extender.

### MEA Recordings

Extracellular recordings were obtained using a Multi Channel Systems (MCS, Reutlingen, Germany) MEA-2100 system and a 120-count multi-electrode array with 100 μm inter-electrode spacing and 30 μm electrode diameter, at a sampling frequency of 20 kHz.

## Author Contributions

D.C.B. conceived of and wrote software and manuscript. K.R.T. provided significant feature guidance and edited manuscript. B.W. provided feature guidance and edited manuscript. K.S.K. and P.K.H. oversaw implementation details and edited manuscript.

## Information Sharing Agreement

MEA Viewer is licensed under the General Public License version 3, GPLv3, and is available at http://github.com/dbridges/mea-tools.

## Acknowledgements

This research was sponsored by the U.S. Army Research Laboratory and Defense Advanced Research Projects Agency under Cooperative Agreement Number W911NF-15-2-0056. The views, opinions, and/or findings contained in this material are those of the authors and should not be interpreted as representing the official views or policies of the Department of Defense or the U.S. Government. Additional support was also provided by the California NanoSystems Institute (CNSI).

## Supplementary Material

### Video

A video displaying MEA Viewer in action is available: https://vimeo.com/143168058

### Spike CSV Format

Spike data is stored in plain text comma-separated value (CSV) files which reside in the same directory as their associated HDF5 recordings and contain the same filename but with a .csv extension instead of .h5. Each file has a single header line detailing 5 columns: “electrode”, “time”, “amplitude”, “threshold”, “conductance”. Each row thereafter specifies a spike event where the electrode column contains the electrode ID, typically in the format A6.0 where A6 is the electrode identifier and 0 specifies the sorted spike group number on that electrode. A spike group number of −1 specifies spikes which could not be properly sorted. The time column specifies the time in seconds that the event occurred in the recording. The amplitude column specifies the amplitude of the event in uV. The threshold column specifies the spike detection threshold used for that channel in uV. The conductance column specifies *True* if this spike event is considered a redundant event due to it being part of a propagation signal, or *False* if it is not. A sample of the data format is given below:

electrode,time,amplitude,threshold,conductance

e12.0,48.861934661865234,-88.80604553222656,-36.17005157470703,False

e12.0,49.07347106933594,-74.42280578613281,-36.17005157470703,False

e12.0,49.107704162597656,-69.9498519897461,-36.17005157470703,True

e12.0,49.15875244140625,-74.18529510498047,-36.17005157470703,False

